# Genetic analyses link the conserved inner membrane protein CvpA to the σ^E^ extracytoplasmic stress response

**DOI:** 10.1101/2020.09.25.314492

**Authors:** Alyson R. Warr, Rachel T. Giorgio, Matthew K. Waldor

## Abstract

The function of *cvpA*, a bacterial gene predicted to encode an inner membrane protein, is largely unknown. Early studies in *E. coli* linked *cvpA* to Colicin V secretion and recent work revealed that it is required for robust intestinal colonization by diverse enteric pathogens. In enterohemorrhagic *E. coli* (EHEC), *cvpA* is required for resistance to the bile salt deoxycholate (DOC). Here, we carried out genome-scale transposon-insertion mutagenesis and spontaneous suppressor analysis to uncover *cvpA’s* genetic interactions and identify common pathways that rescue the sensitivity of a Δ*cvpA* EHEC mutant to DOC. Collectively, these screens led to the hypothesis that the Δ*cvpA* mutant is impaired in its capacity to activate the σ^E^-mediated stress response. This idea was supported by showing that mutations that activate σ^E^, either indirectly or through its direct overexpression, can restore the Δ*cvpA* mutant’s resistance to DOC. Analysis of the distribution of CvpA homologs revealed that this inner membrane protein is conserved across bacterial phyla, in both enteric and non-enteric bacteria that are not exposed to bile. Together, our findings suggest that CvpA may function in triggering activation of the σ^E^ stress response pathway in response to DOC as well as additional stimuli.

**Importance:** Several enteric pathogens, including Enterohemorrhagic E. coli (EHEC), require *cvpA* to robustly colonize the intestine. This inner membrane is also important for secretion of a colicin and EHEC resistance to the bile salt deoxycholate, but its function is unknown. Genetic analyses carried out here suggest that *cvpA* is required to trigger the σE stress response pathway in response to deoxycholate. Since CvpA is conserved across diverse bacterial phyla, we propose that this inner membrane protein is important for activation of this stress response pathway in response to diverse perturbations of the cell envelope.

## Introduction

Enteric pathogens encounter a broad range of host-derived stressors during their transit through the gastrointestinal (GI) tract. These pathogens enter the GI tract through consumption of contaminated food or water, and the diverse challenges they face in the host environment include substantial fluctuations in osmolarity, oxygen concentration, pH, and nutrient availability, as well as mechanical shear force from peristalsis and the bactericidal activities of antimicrobial peptides and bile salts. Pathogens must also compete with the microbiota and face the threat of immune cells and effectors (1, 2). To survive and successfully colonize the host, enteric pathogens must rapidly sense and respond to these stressors.

Bile in particular poses a complex threat to enteric pathogens. Bile is an aqueous solution of bile salts, bilirubin, fats, and inorganic salts produced by the liver and secreted into the intestine, where it plays a critical role in digestion of fats (3, 4). Bile salts are potent antimicrobial compounds that can damage diverse components of the bacterial cell. Bile salts are thought to enter the bacterial cell by both passive diffusion and through outer membrane proteins (OMPs) such as porins (5, 6). Enteric pathogens employ a variety of strategies to combat the perturbations caused by bile salts, including utilizing efflux pumps to remove bile salts from the cell cytosol, and expressing chaperones and proteases to ameliorate protein folding stress (7–9).

To coordinate responses to bile, bacteria utilize a host of global stress-response programs. For example, bile-induced DNA damage activates the expression of the SOS regulon, a large suite of genes that assist in repairing and responding to DNA damage (10–12). Bile also triggers the activation of alternate sigma factors, which enable RNA polymerase (RNAP) to selectively transcribe sets of genes in response to environmental conditions. During normal growth conditions, RNAP associates with the housekeeping sigma subunit, RpoD (σ^70^). The cellular stresses provoked by bile promote utilization of the alternative stress response sigma factor RpoS (σ^S^), which governs expression of a large number (>500) of stress-response genes (13–15). RpoE (σ^E^), a sigma factor which mediates the response to ‘extracytoplasmic’ stress, such as misfolded proteins in the periplasmic space or disruptions in LPS biosynthesis, may also be important for resistance to the membrane-damaging effects of bile (16, 17). σ^E^ function intersects with a variety of other cell envelope stress response pathways in Gram-negative bacteria such as the two-component systems CpxRA and BaeSR; together, these pathways create a complex regulatory network mediating bacterial envelope homeostasis (16, 18).

The gene *cvpA* was identified in screens for genes required for intestinal colonization in several enteric pathogens, including *Vibrio cholerae*, *Vibrio parahaemolyticus*, and *Salmonella* Typhimurium (19–21), but the mechanism(s) that account for the colonization defects have not been characterized. We recently found that CvpA is required for the foodborne intestinal pathogen enterohemorrhagic *Escherichia coli* O157:H7 (EHEC) to optimally colonize the colon of infant rabbits; moreover, EHEC *cvpA* deletion mutants were highly sensitive to bile, and in particular the bile salt deoxycholate (DOC) (22). EHEC causes hemorrhagic colitis and in some cases the severe complication hemolytic uremic syndrome, a condition in which damage to blood vessels of the kidney microvasculature leads to renal failure. Previous studies have led to the idea that EHEC uses bile as a cue to induce expression of virulence genes, flagella, and genes promoting survival *in vivo*, including the AcrAB efflux pump which enables removal of bile salts from the cell interior (12, 23–29), but the role of CvpA in contributing to bile resistance in EHEC is unknown.

*cvpA* was originally described as a gene required for the production of Colicin V (ColV), a small peptide antibiotic produced by some strains of pathogenic *E. coli* (30). Although the ColV secretion system was characterized (31), the specific role for CvpA in this mechanism was not identified. Here, we provide evidence that CvpA likely provides EHEC resistance to DOC by modulating the σ^E^ stress response pathway. We also show that CvpA is widely conserved across diverse bacterial phyla and propose that it may function to coordinate stress responses to perturbations in ion homeostasis.

## Results

### CvpA localizes to the EHEC cell periphery

CvpA bears structural similarity to the inner membrane mechanosensitive ion channels MscS and MscL (HHPred, Phyre2) (22), and whole-proteome localization analysis in *E. coli* assigned CvpA to the inner membrane (32). Furthermore, several protein modeling algorithms (PSLPred, HHPred, Phobius, Phyre2, OCTOPUS) (22) predict that CvpA is a 4-pass inner membrane protein with a periplasmic C-terminus (Fig 1A). To experimentally verify these assignments, we generated two plasmids that inducibly express sfGFP-CvpA fusions, with sfGFP on either the N- or C-terminus, enabling visualization of the localization of CvpA in EHEC. Only the sfGFP fusion to the N-terminus of CvpA yielded a construct that complemented the growth defect of an EHEC strain lacking the *cvpA* gene (Δ*cvpA*) on plates containing DOC, implying this construct contains a functional CvpA protein (Fig 1B). Induction of this fusion protein revealed a clear signal that outlined the EHEC cell membrane in a similar pattern as the membrane stain FM4-64 (Fig 1CD). These data support the predicted localization of CvpA, and suggest that the availability of its periplasmic C-terminus may be necessary for function.

**Figure 1:**
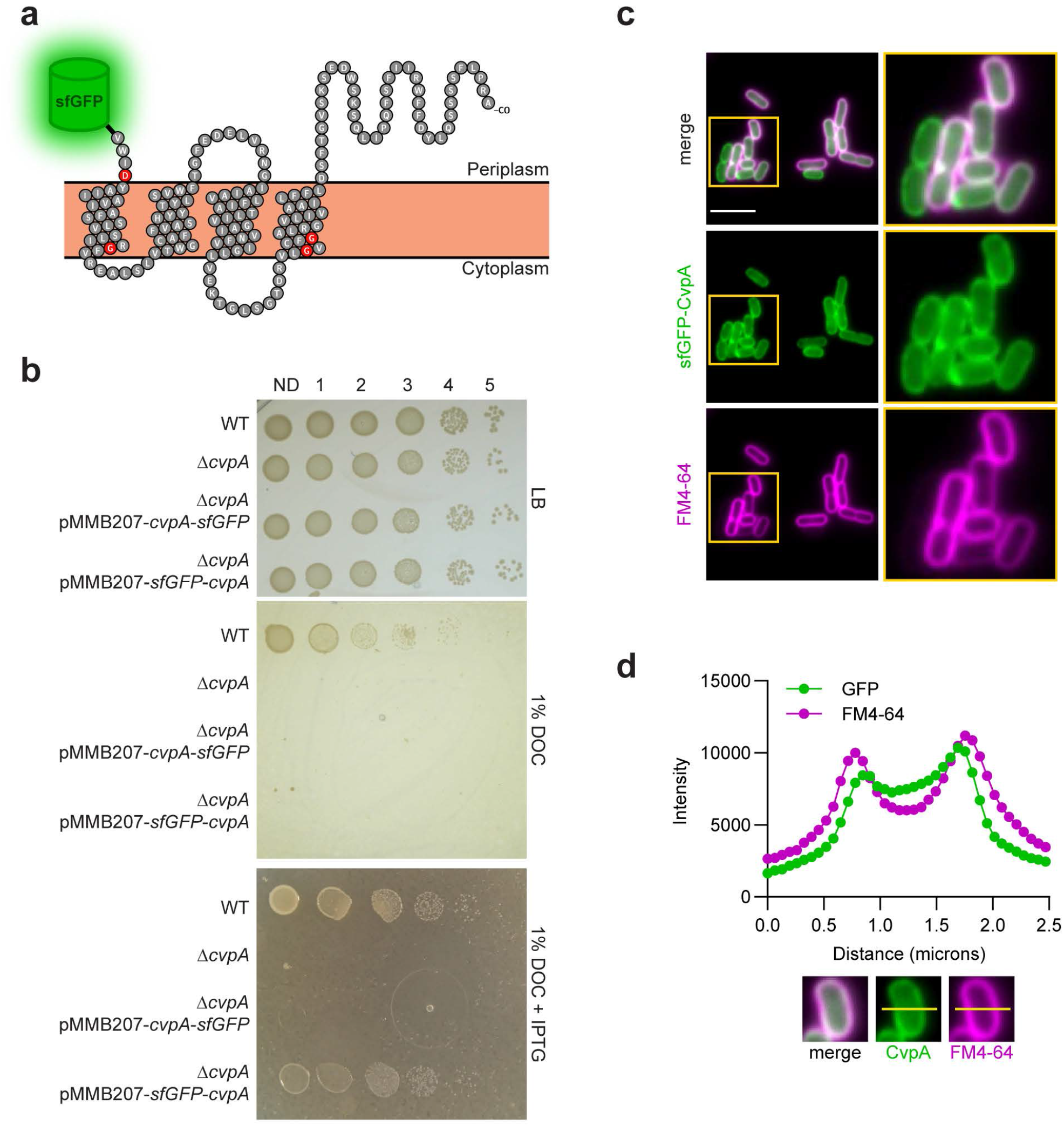
CvpA localizes to the EHEC cell periphery. a) Schematic of the predicted CvpA topology (adapted from (22)), with location of N-terminal sfGFP fusion shown. Red residues are nearly 100% conserved across bacterial phyla. b) Dilution series of WT, Δ*cvpA* mutant, and Δ*cvpA* mutant with an IPTG-inducible *sfGFP*-*cvpA* fusion complementation plasmid plated on LB, LB 1% deoxycholate (DOC), and LB 1% DOC 1 mM IPTG. c) Micrographs of an N-terminal sfGFP-CvpA fusion protein expressed in EHEC and stained with the FM4-64 membrane dye. Scale bar indicates 5 μM. d) Horizontal line scan of GFP and FM4-64 signal.

### EHEC *cvpA* mutants do not secrete functional ColV and are sensitive to the bile salt DOC

A previous report suggested that *E. coli* CvpA is required for the export of the peptide antibiotic Colicin V (ColV) (30). We sought to reproduce this observation in EHEC. The entire ColV-production and immunity operon and endogenous promoters derived from the naturally occurring ColV plasmid, pColV-K30, was inserted into the cloning vector pBR322, yielding pBR322-ColV. Although both WT (EDL933) and Δ*cvpA* EHEC were transformable with pBR322-ColV, only the WT strain grew robustly in LB broth while carrying pBR322-ColV. In contrast, the Δ*cvpA* mutant elongated and lysed (Fig S1A). The abilities of these strains to secrete functional ColV was evaluated by testing whether they create a zone of growth inhibition on a sensitive indicator strain. The WT strain carrying pBR322-ColV killed the indicator strain, creating a distinct zone of clearing (Fig S1B), whereas the Δ*cvpA* strain failed to establish such a zone (Fig S1B). These data are consistent with previous reports (30) and suggest that Δ*cvpA* mutants are unable to secrete functional ColV and may be sensitive to self-intoxication.

Our previous work revealed that CvpA is required for EHEC to robustly colonize the infant rabbit colon, and that Δ*cvpA* mutants are sensitive to the bile salt DOC but not to the bile salt cholate (CHO) (22). Bile salts like DOC are thought to cause a wide range of deleterious effects in bacteria, including damage to the cell envelope and DNA, and widespread protein misfolding and redox stress (5, 6). The exact mechanisms through which bile salts mediate these diverse deleterious effects are not completely understood (5, 6). We attempted to determine which aspect of DOC toxicity was most damaging to Δ*cvpA* mutants by testing a variety of agents that cause individual aspects of bile stress, including the membrane-damaging detergent SDS, DNA damaging antibiotics, cell wall perturbing antibiotics, protein synthesis disrupting antibiotics, and the redox-stress inducing compounds diamide and hydrogen peroxide (22) (Table 1). However, the Δ*cvpA* mutant was not more sensitive than the WT strain to any of these agents. These observations argue against the idea that the sensitivity of the Δ*cvpA* mutant to DOC results from a general cell envelope defect, and instead suggest that DOC toxicity (and potentially the toxicity of ColV accumulation) reflects a more specific defect in cells lacking *cvpA*. Below, we moved forward with DOC as a tool to probe the function of CvpA.

**Table 1:**
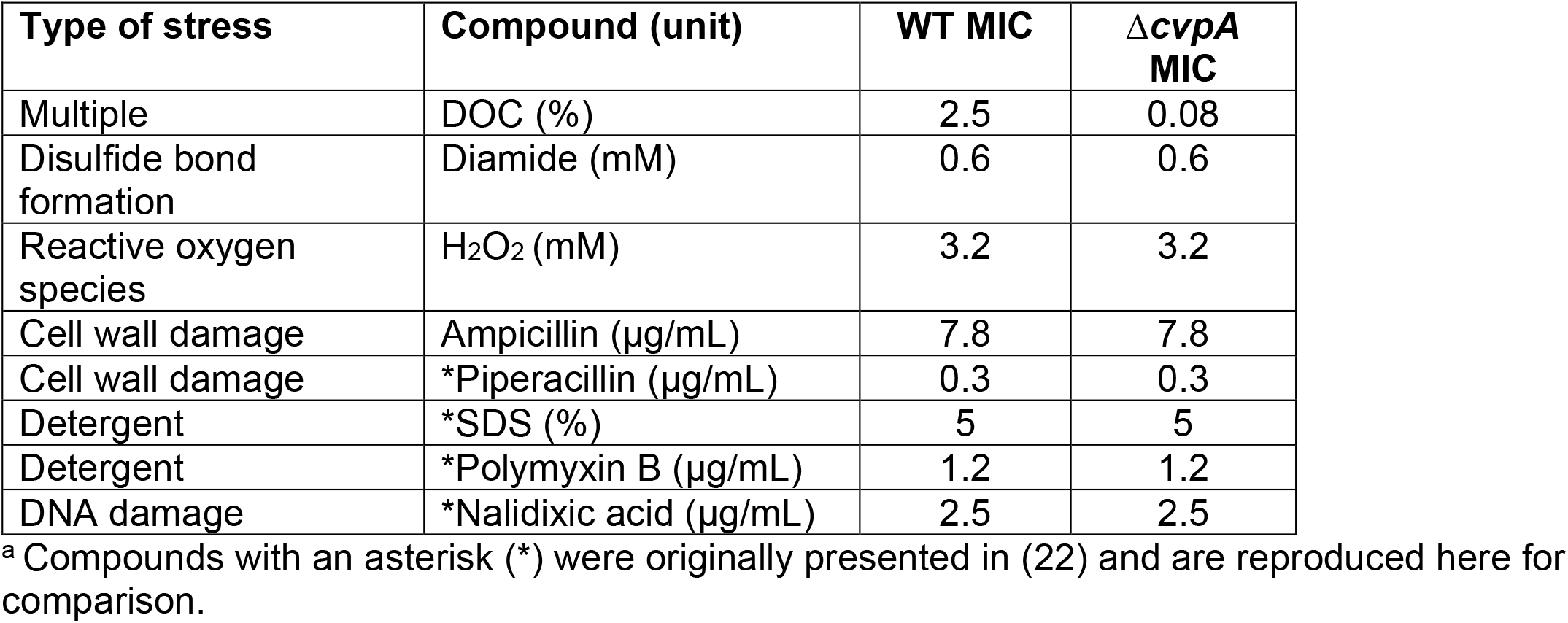
MIC for various compounds in WT and Δ*cvpA* EHEC^a^.

| Type of stress | Compound (unit) | WT MIC | $\Delta cvpA$ MIC |
| --- | --- | --- | --- |
| Multiple | DOC (%) | 2.5 | 0.08 |
| Disulfide bond formation | Diamide (mM) | 0.6 | 0.6 |
| Reactive oxygen species | H<sub>2</sub>O<sub>2</sub> (mM) | 3.2 | 3.2 |
| Cell wall damage | Ampicillin (μg/mL) | 7.8 | 7.8 |
| Cell wall damage | *Piperacillin (μg/mL) | 0.3 | 0.3 |
| Detergent | *SDS (%) | 5 | 5 |
| Detergent | *Polymyxin B (μg/mL) | 1.2 | 1.2 |
| DNA damage | *Nalidixic acid (μg/mL) | 2.5 | 2.5 |

<sup>a</sup> Compounds with an asterisk (\*) were originally presented in (22) and are reproduced here for comparison.

### Genetic screens link CvpA to the σ^E^ response

To further assess CvpA’s function, we carried out three genetic screens in EHEC to define *cvpA’s* network of genetic interactions. Two of the screens relied on transposon-insertion mutagenesis, and one used spontaneous suppressor analysis. Similar genetic approaches have been fruitful for other membrane proteins (33–36), which are often difficult to probe using biochemical approaches to identify protein-protein interactions.

For the transposon screens, we began by generating high density transposon-insertion mutant libraries in the WT and Δ*cvpA* EHEC strains by mutagenizing the strains with the mariner transposon Himar1, which inserts randomly at TA dinucleotides. The abundance of each transposon-insertion mutant in the two libraries was quantified by deep sequencing the sites of transposon insertion. Genes were binned by the percentage of TA sites disrupted (Fig 2A), and as expected for libraries of high insertion density, this distribution was bimodal. The left-most minor peak represents genes with few to no insertions, and the right-most major peak represents genes with a majority of TA sites disrupted (37). The distribution of transposon insertions in the WT and Δ*cvpA* libraries was similar and displayed complexity enabling high-resolution analysis of transposon-insertion frequency (19) (Fig 2A).

**Figure 2:**
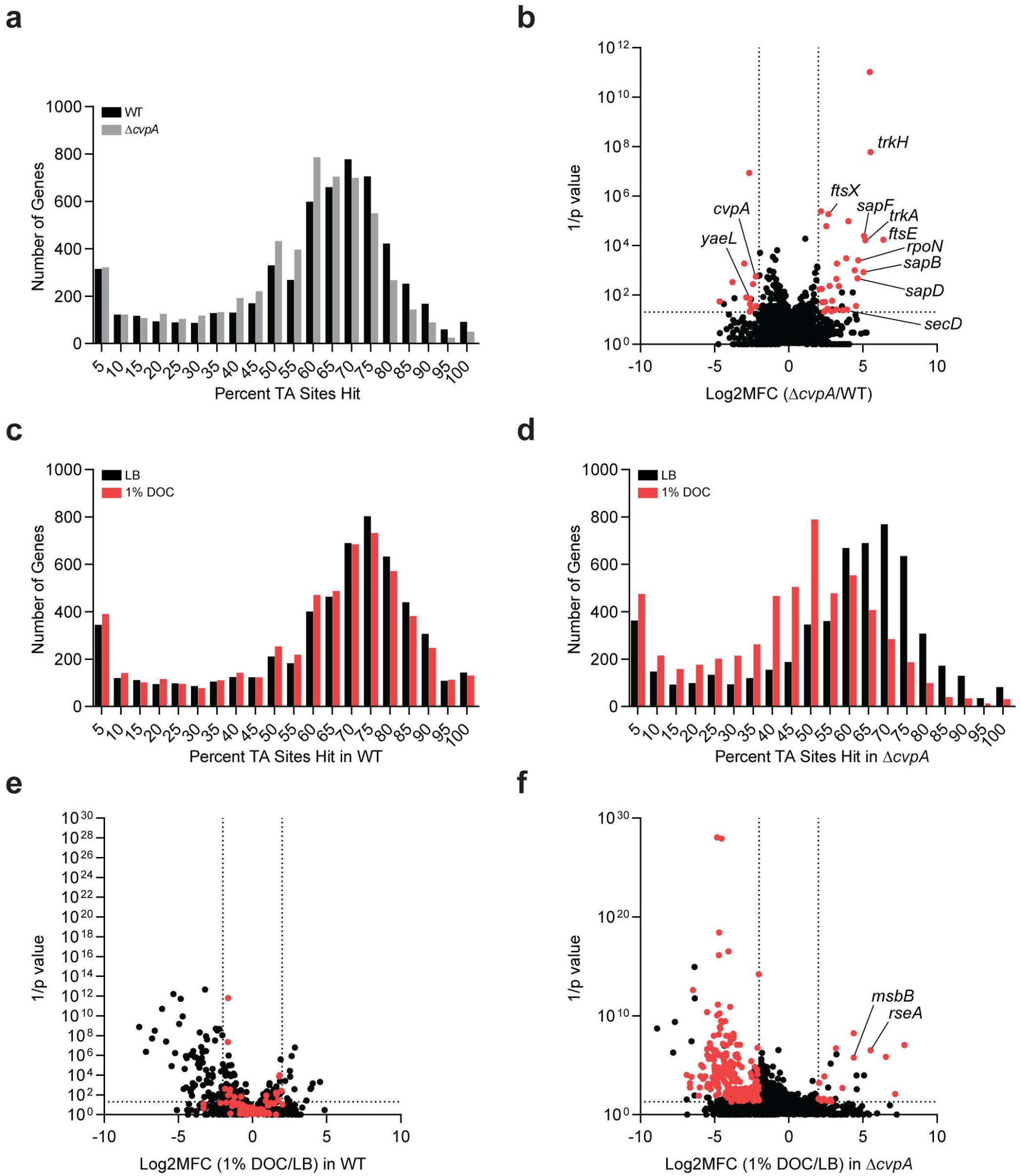
Transposon-insertion sequencing screens reveal CvpA’s genetic interactions. a) Distribution of the percentage of TA sites disrupted for all genes in the library constructed in the WT (black bars) and Δ*cvpA* (gray bars) backgrounds. b) Volcano plot depicting the relative abundance of read counts mapped to individual genes in transposon libraries in WT and Δ*cvpA* mutant backgrounds. The mean log2 fold change (x axis) and Mann-Whitney U p-value (y axis) are shown for each gene. Genes highlighted in red have greater than 5 unique transposon mutants, a log2FC>2 or <−2, and p-value < 0.05. c-d) Distribution of the percentage of TA sites disrupted for all genes in the library constructed in the WT (c) or Δ*cvpA* background (d) after growth in LB (black bars) or 1% DOC (red bars). e-f) Volcano plot depicting the relative abundance of read counts mapped to individual genes in transposon libraries in WT (e) or Δ*cvpA* background (f) in LB vs in 1% DOC. The log2 mean fold change (x axis) and Mann-Whitney U p-value (y axis) are shown for each gene. Genes highlighted in red are those for which mean log2FC >2 or <−2 and p-value < 0.05 in the Δ*cvpA* mutant background but not the WT background.

We first completed a synthetic screen in which we compared the distributions of the transposon insertions across the genome in the WT and Δ*cvpA* libraries. This comparison yields three categories of genes: (1) loci with similar insertion profiles in both backgrounds are deemed “neutral”; (2) loci with fewer insertions in the Δ*cvpA* vs the WT background are considered “synthetic sick”; and (3) loci with more insertions in the Δ*cvpA* compared to the WT background are deemed “synthetic healthy”. This approach has been used to identify genetic interactions in several other organisms (38–41). We identified 32 candidate synthetic healthy genes, for which transposon-insertion mutants were overrepresented at least 4-fold in the Δ*cvpA* mutant background compared to the WT (Fig 2B, Sup Table 1). Several of these genes were linked to ion transport systems, including *trkH* and *trkA*, which are components of the cellular potassium uptake system, and three genes of the *sap* transporter complex (*sapB, sapD*, and *sapF)*. SapD is also known to interact with the Trk potassium transport system (42). Several chaperones, including *secB*, the alternative sigma factor *rpoN*, and several cell shape/septal structure genes, including *ftsEX*, were also on the list of synthetic healthy genes. 14 candidate synthetic sick genes, which were underrepresented at least 4-fold in the Δ*cvpA* mutant background compared to the WT (Fig 2B, Sup Table 1), were also identified. These genes included *yaeL*, a protease that works with DegS to activate σ^E^ by degrading the anti-sigma factor RseA. Collectively, these results suggest that inactivating certain ion transport processes bolster the fitness of the Δ*cvpA* strain, while impairing the cell’s ability to activate the σ^E^-mediated stress response is particularly deleterious in the Δ*cvpA* background. Interestingly, there is evidence that cellular potassium levels can regulate promoter selectivity of sigma factors (43).

A second transposon screen was designed to identify insertion mutations in genes that suppress the DOC sensitivity phenotype in the Δ*cvpA* mutant. The WT and Δ*cvpA* transposon-insertion mutant libraries were screened on plates containing either LB or LB supplemented with 1% DOC. As above, after enumerating the abundance of transposon-insertion mutants with sequencing, genes were binned according to the percentage of disrupted TA sites (Fig 2CD). The distribution of disrupted TA sites per gene in the WT library was not markedly different between the LB and DOC selection, as expected because a majority of mutants do not have a phenotype in DOC (Fig 2C). However, the distribution in the Δ*cvpA* library was different between the two conditions; there was a leftward shift in the distribution of insertions after the Δ*cvpA* library was exposed to DOC compared to LB (Fig 2D). This leftward shift is indicative of a bottleneck, a stochastic, genotype-independent constriction of population size, and reflects the profound DOC sensitivity of the Δ*cvpA* mutant.

To identify mutants in both libraries which have markedly better or worse phenotypes than expected in DOC, we used Con-ARTIST, an analytical pipeline that mitigates the effects of bottlenecks to identify differentially abundant mutants (19). For each library, the abundance of each mutant was compared between the LB and DOC condition to identify three categories of genes: (1) loci with similar insertion profiles in both LB and DOC are “neutral”; (2) loci with fewer insertions in the DOC condition than expected when compared to LB are “depleted”; and (3) loci with more insertions in the DOC condition than expected when compared to LB are “enriched” (Fig 2EF). In the Δ*cvpA* background, we identified 233 ‘depleted’ genes in which mutations are expected to increase baseline DOC sensitivity, and 26 ‘enriched’ genes in which mutations are expected to suppress DOC sensitivity (Fig 2F). 19 of these 26 genes did not show differential abundance in DOC vs. LB comparison in the WT background and were classified as ‘neutral’ (Fig 2EF). These genes are of particular interest as candidate suppressors, as they have *cvpA*-specific DOC phenotypes. Many of these genes were linked to stress response pathways and cell envelope integrity, such as the anti-sigma factor *rseA*, the σ^E^ regulated periplasmic chaperone *skp*, the two-component system *basSR*, the redox genes *sodC* and *cyoC*, the disulfide/redox bond formation genes *dsbA* and *dsbB*, and the cell membrane synthesis genes *msbB, wcaM*, and *rfaC*. Mutations in several of these genes are known to increase the level of σ^E^ activity (44–46).

In addition to transposon-based screens, we also characterized spontaneous suppressors of DOC sensitivity in the Δ*cvpA* mutant. In this screen, 2 mL of OD 0.5 Δ*cvpA* broth culture were plated on large plates and 16 putative suppressor colonies were picked after 24 hours. After re-patching on 1% DOC, which confirmed their resistance, whole-genome sequencing of the suppressors was carried out and these genomes sequences were compared to that of the Δ*cvpA* parent strain to identify putative suppressor mutations. SNPs or deletions that were not present in Δ*cvpA* parent strain were found in 14 of the 16 strains. Notably 12 of the 14 strains had mutations in genes linked to LPS biogenesis, including several genes in the lipopolysaccharide transport (LPT) system (*lptB, lptE, lptF, lptG*) and an enzyme required for lipid A synthesis, *msbB;* genes related to redox including *nuoH* and *cyoDC* were also identified (Table 2). Interestingly, *msbB* was also identified as a candidate suppressor in the transposon DOC screen above. LPT genes are essential, and thus not highly represented in our transposon libraries. Although not statistically robust, we analyzed the transposon DOC screen data for the LPT genes with at least 1 insertion mutant and found that *lptCDFG* had large, positive fold-changes in 1% DOC vs LB in the Δ*cvpA* background and not WT (Sup Table 2), supporting the findings of the spontaneous suppressor screen.

**Table 2:**
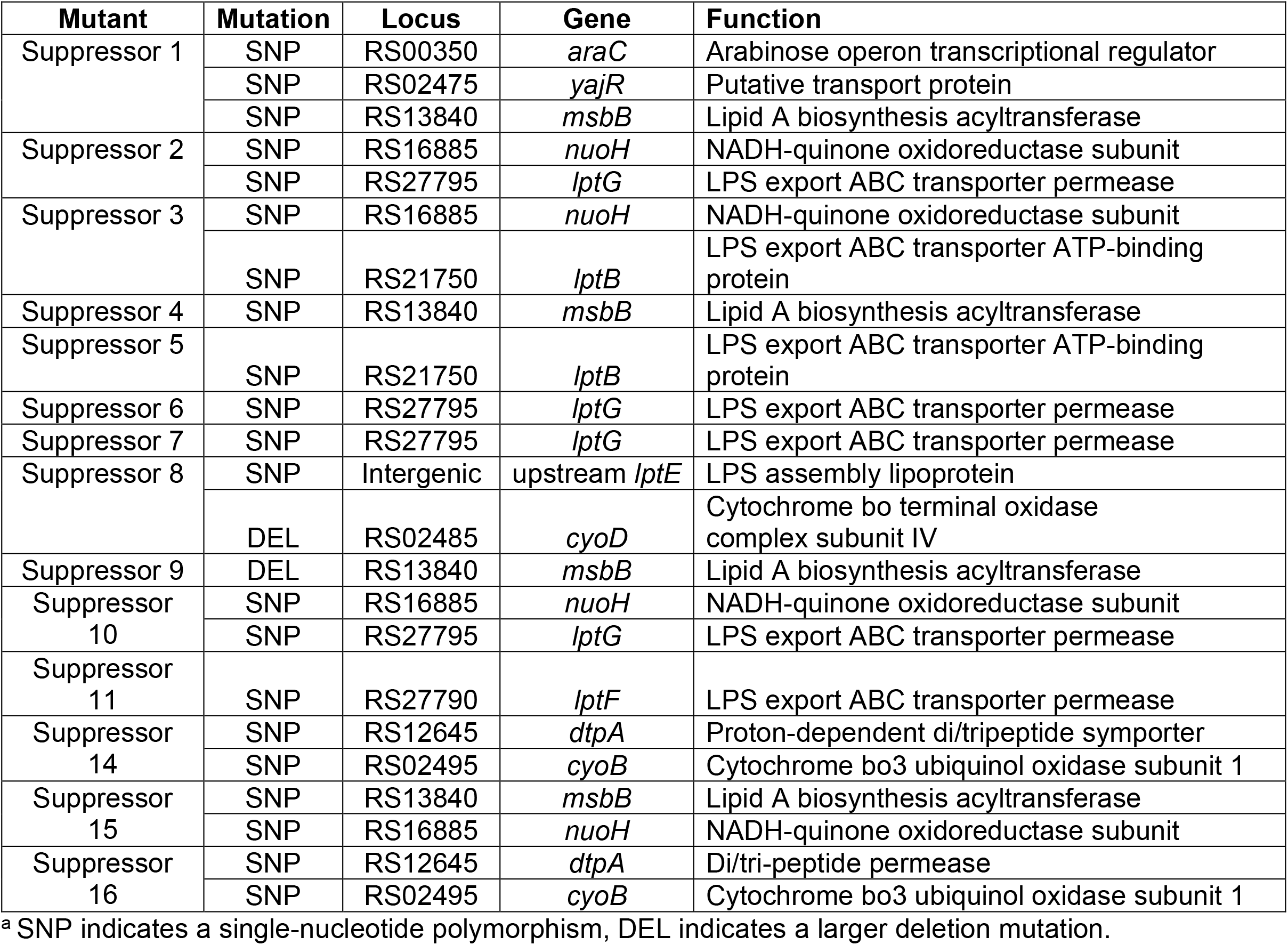
Mutations in suppressors of Δ*cvpA* DOC sensitivity^a^.

### Increased σ^E^ activity rescues the sensitivity of the Δ*cvpA* mutant to DOC

Several independent pieces of evidence connecting *cvpA* to the σ^E^ response were found in the 3 genetic screens described above. σ^E^ is normally sequestered at the inner membrane by association with an anti-sigma factor, RseA. When the appropriate stimuli is detected, proteases YaeL and DegS sequentially cleave RseA, liberating σ^E^ to activate expression of ~100 genes in response (47, 48). A variety of perturbations to the cell envelope can trigger this pathway, including misfolded OMPs, disruptions in LPS biosynthesis leading to precursor accumulation, and changes in cellular redox state (49, 44–46). Notably, the genetic screens revealed that the Δ*cvpA* mutant is more sensitive than the WT to mutations in *yaeL* and the suppressor screens uncovered a variety of mutations, including in *rseA*, *lptBCDEFG*, and *msbB*, which likely trigger the σ^E^ response, that appeared to rescue the sensitivity of Δ*cvpA* to DOC. Inactivating mutations in *rseA* increase the basal activity of σ^E^ by preventing its membrane sequestration, and mutations in LPS biosynthesis genes *msbB* and the LPS transport system trigger σ^E^ through accumulation of LPS precursors (44, 45).

To test the linkage between *cvpA* and σ^E^, deletion mutants of *rseA and msbB* in the WT and Δ*cvpA* backgrounds were created. These deletions both restored the resistance of the Δ*cvpA* mutant to 1% DOC to WT levels (Fig 3A). Similarly, direct overexpression of *rpoE* from a plasmid rescued the sensitivity of the Δ*cvpA* mutant to DOC (Fig 3B). These data suggest that mutations that activate σ^E^, either indirectly or through its direct overexpression, can restore the Δ*cvpA* mutant’s resistance to DOC, implying that the mutant may be otherwise unable to activate σ^E^ in response to DOC.

**Figure 3:**
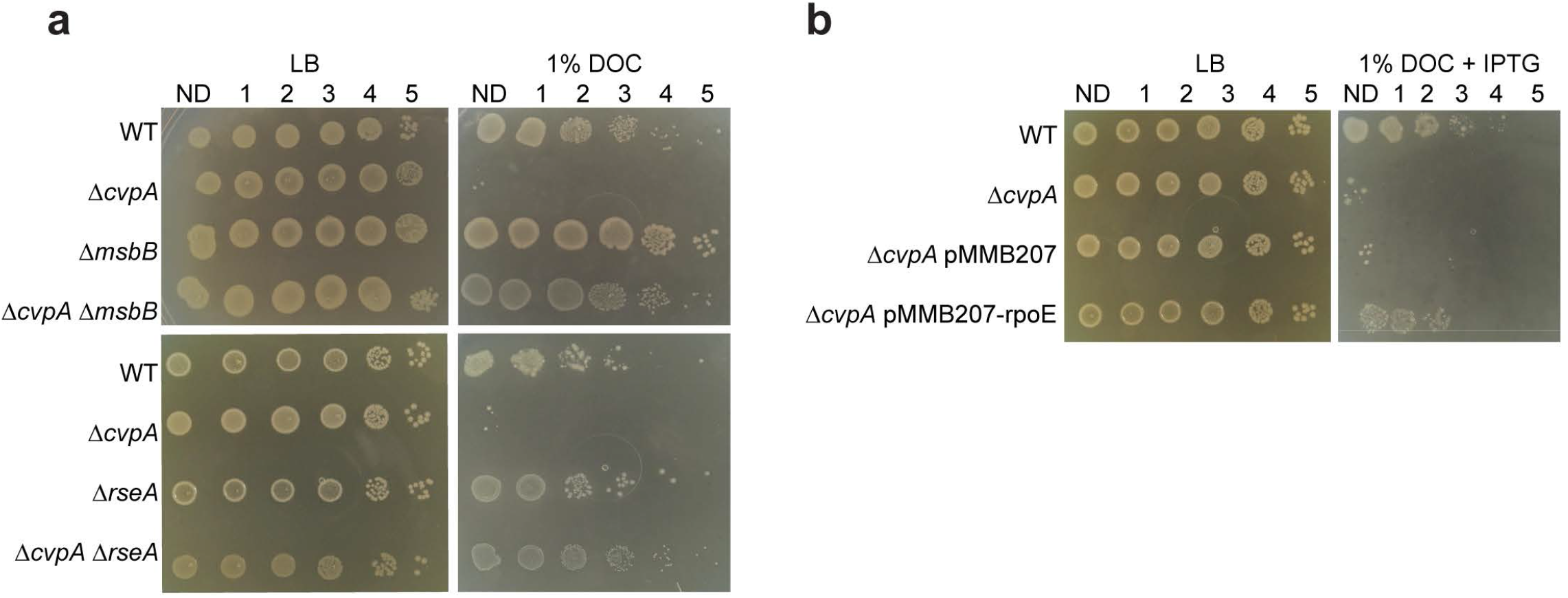
Activation of RpoE rescues the Δ*cvpA* mutant’s sensitivity to DOC. a) Dilution series of WT, Δ*cvpA*, Δ*cvpA* Δ*msbB* and Δ*cvpA* Δ*rseA* plated on LB or LB 1% deoxycholate (DOC). b) Dilution series of WT, Δ<u>*cvpA*</u>, Δ*cvpA* transformed with empty expression vector pMMB207, and Δ*cvpA* transformed with pMMB207 carrying IPTG inducible *rpoE* plated on LB and LB 1% DOC with 0.1 mM IPTG.

### CvpA is highly conserved across bacterial phyla

The phylogeny tool Annotree (50) was used to analyze the distribution of CvpA homologs across more than 27,000 bacterial genomes. CvpA homologs (≥30% amino acid identity, ≥70% alignment) were found in 9,830 genomes distributed across 55 phyla (Fig 4A, Sup Table 3). Homologs are present in the majority of represented genomes in the Campylobacterota (236/254, 93%), Deferribacterota (13/14, 93%), and Proteobacteria (6923/7630, 91%) phyla (Fig 4B). In contrast, only 2 of 3118 strains in Actinobacteria have a CvpA homolog (0.06%) (Fig 4B).

**Figure 4:**
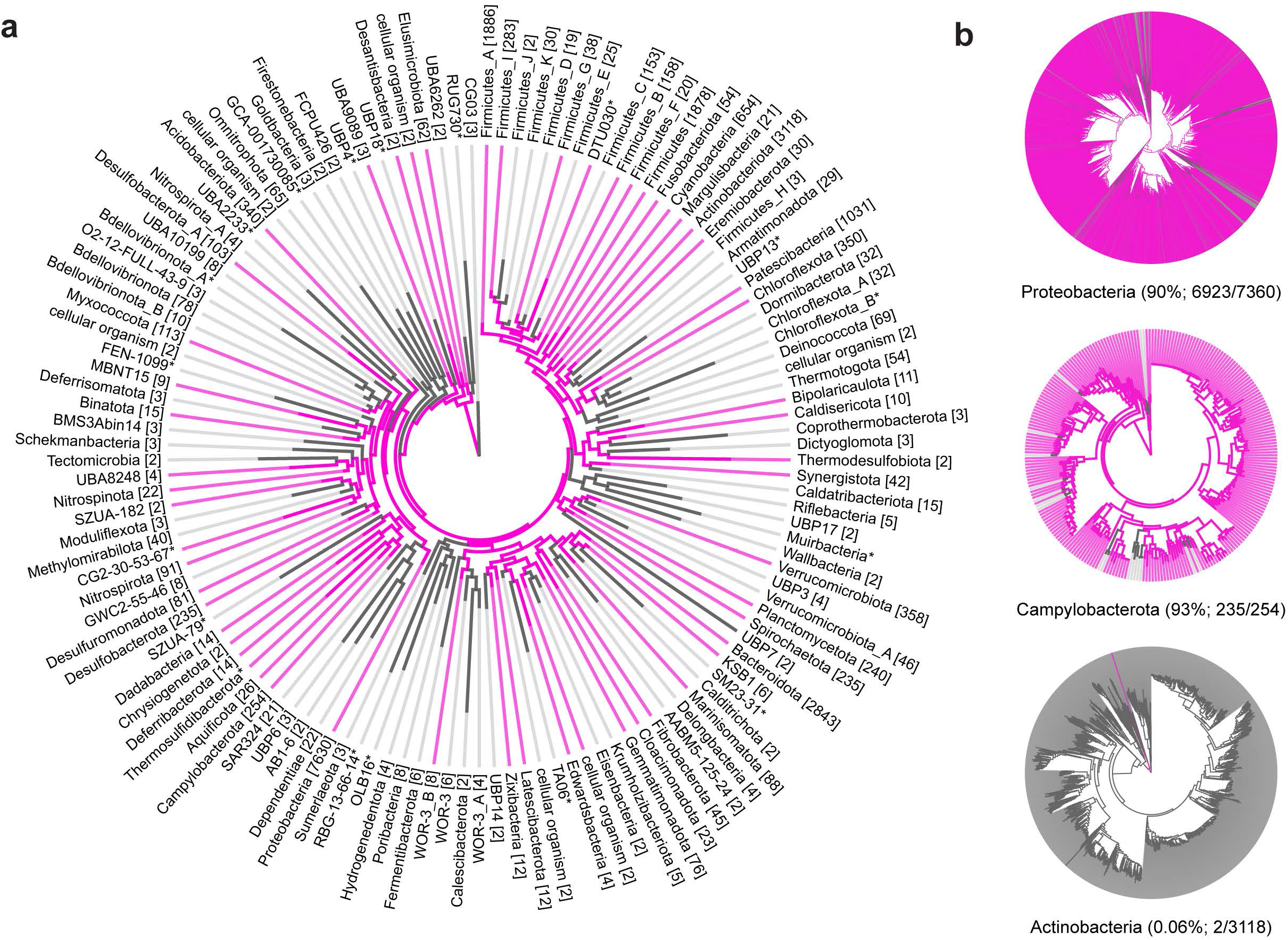
CvpA is highly conserved across many bacterial phyla. a) Phylogeny of CvpA distribution across bacterial phyla. Pink lines represent phyla where at least one species has a CvpA homolog of at least 30% amino acid identity and at least 70% alignment to *E. coli* CvpA. The numbers shown in brackets denote the number of individual genomes in that phylum. b) Right, phyla with varying degrees of CvpA conservation at the species level. Pink lines represent species that contain a CvpA homolog. Numbers in parenthesis after phylum name indicate the percentage of species in that phylum with a CvpA homolog.

Sequence alignment of the primary amino acid sequences of 20 CvpA homologs from bacterial species from diverse phyla revealed several highly conserved residues, including an aspartate and three glycines with nearly 100% conservation (Fig S2, highlighted in red in Fig 1A). The motif GXXXG, common in transmembrane segments and indicative of helix-helix interactions (51), is also present. All 20 CvpA homologs are predicted to localize to the inner membrane by PSLPred. All homologs share greater similarity in alignments of their respective N-terminal regions, and have more variable lengths in their final periplasmic segments, which could reflect different functions or binding partners at the C-terminus.

This conservation across diverse phyla highlights that the function of CvpA is not strictly related to bile. While homologs are present in enteric pathogenic and commensal strains, such as *Campylobacter jejuni, Yersinia enterolitica*, *Fibrobacter succinogenes*, and *Bacteroides fragilis*, many strains with a CvpA homolog inhabit non-intestinal niches. These strains include non-enteric pathogens such as *Neisseria meningitidis*, plant symbionts such as *Rhizobium tropici*, and non-pathogenic environmental isolates from diverse environments such as tundra soil (*Granulicella tundricola*), hydrothermal vents (*Thermovibrio ammonificans*), and marine environments (*Spirochaeta cellobiosiphila*).

## Discussion

Since the original report describing *cvpA* as a gene required for production of the plasmid-borne ColV peptide from *E. coli* more than 30 years ago (30), very little progress has been made determining the function of this widely conserved inner membrane protein. More recently, *cvpA* has been found to be important for intestinal colonization of *Vibrio cholerae*, *V*. *parahaemolyticus*, and *Salmonella* Typhimurium and EHEC in studies based on *in vivo* transposon screens (19–22). In EHEC, we found that a Δ*cvpA* mutant was highly sensitive to DOC (22), a bile component, thus presenting a possible explanation for *cvpA*’s role in the colonization of distinct niches in the small and large intestine by diverse enteric pathogens. Here, we used genetic screens to further our understanding of *cvpA* function. These screens led to the hypothesis that *cvpA* is an upstream component of the σ^E^ ‘extracytoplasmic’ stress response pathway that responds to DOC and other stimuli, such as ColV secretion.

A variety of stimuli can lead to the activation of the σ^E^ response. Misfolded outer membrane proteins, disruptions in LPS biogenesis, and perturbations in redox state have all been linked to increased σ^E^ activity (49, 44–46). Mutations in the regulators of these pathways, for example inactivating *rseA*, the anti-sigma factor which controls σ^E^ activity (49), *msbB*, an enzyme required for the biosynthesis of lipid A (44), or the redox enzymes *cyoC, sodC*, and *dsbAB* (46) can also trigger the activation of σ^E^ in the absence of external stimuli. Conversely, inactivating mutations in *degS* and *yaeL*, the proteases required to release sequestered σ^E^ from the membrane, inhibit σ^E^ activation (52–54). Nearly all of the σ^E^-interacting genes listed above were identified as potential modulators of the fitness Δ*cvpA* mutant in DOC (Fig 2, Table 2). These data suggest that in response to DOC, CvpA may be required to activate the σ^E^ response. In a Δ*cvpA* mutant, this pathway is unable to respond to deleterious stimuli, and additional σ^E^ - activating mutations (deletions in *rseA, msbB, lpt* genes, direct overexpression of *rpoE*) are required to bypass this block (Table 2, Fig 3). Mutations in *cyoC, sodC*, and *dsbAB* were unable to suppress the sensitivity of Δ*cvpA* DOC mutations, despite these loci having strong phenotypes in the DOC transposon-insertion screen. It is possible that under the conditions of our *in vitro* validation experiments, deletions in these enzymes are not sufficient to rescue DOC sensitivity.

The conservation of CvpA across diverse bacterial phyla, including in species which are never exposed to bile, suggests that CvpA function is not restricted to responding to bile stress (Fig 4). Bile can impair several processes in the bacterial cell, including inflicting damage to the cell envelope and DNA, and generating protein folding stress and alterations in redox state (5, 6). We were unable to isolate which individual aspect of DOC stress kills Δ*cvpA* mutants (Table 1), raising the possibility that either these deleterious effects need to occur together, or that bile impairs additional processes that were not tested. One possibility is that bile, and DOC in particular, perturbs ion homeostasis, which could in turn disrupt membrane potential. The increased fitness of insertions in several genes associated with K^+^ transport (Fig 2B) links *cvpA* and ion transport processes and supports this idea. Furthermore, in eukaryotic cells, DOC exposure leads to changes in mitochondrial membrane potential (55). Interestingly, Δ*cvpA* mutants are not sensitive to the very similar bile salt cholate (CHO) (22), which may reflect a difference in the ability of these salts to move through membranes (56) or the increased hydrophobicity of DOC (57).

Given the similarity of CvpA to the mechanosensitive solute transporters MscS and MscL and the numerous genes related to ion transport that were identified in the synthetic lethal screen (Fig 2B), it is tempting to speculate that CvpA is itself an ion channel that monitors and promotes membrane potential homeostasis. Notably, Colicin V mediates its bactericidal action by disrupting the membrane potential of the inner membrane (58, 59). If Δ*cvpA* mutants have heightened sensitivity to perturbations of membrane potential, this could explain why they cannot produce functional Colicin V. Maintenance of ion homeostasis and membrane potential is a critical capability for all bacterial species. DOC may represent a stimulus that leads to perturbation of ion levels; other stimuli that lead to similar perturbations are likely encountered in the disparate niches inhabited by the diverse bacterial species encoding CvpA homologs.

Finally, our observations suggest that the C-terminus of CvpA is required for its function. A fusion of sfGFP to this end of the protein was not functional. The C-terminal region of CvpA varies in length and sequence between species. Albeit speculative, we propose that this region of the protein may be critical for binding to periplasmic partner(s) or sensing/responding to periplasmic stimuli. Clearly, additional biochemical studies to elucidate the mechanistic bases of CvpA function are warranted.

## Materials and Methods

### Bacterial strains and growth conditions

Bacterial strains were cultured in LB medium or on LB agar plates at 37°C. Antibiotics and supplements were used at the following concentrations: 20 μg/mL chloramphenicol (Cm), 50 μg/mL kanamycin (Km), 10 μg/mL gentamicin (Gm), 50 μg/mL carbenicillin (Cb), 0.3 mM diaminopimelic acid (DAP), 1% deoxycholate (DOC), 0.1 mM Isopropyl β-d-1-thiogalactopyranoside (IPTG).

A gentamicin-resistant mutant of *E. coli* O157:H7 strain EDL933 (Δ*lacI*::*aacC1*) (22) was used in all experiments in this study as the wild-type (WT). All mutants were constructed in this strain background using standard allelic exchange techniques (60) with the pTOX plasmid system (61) or lambda-red recombineering (62).

Colicin V production and immunity operons and endogenous promoters were cloned from plasmid pColV-K30:Tn10 (gift from Dr. Roberto Kolter) into pBR322 (yielding ‘pBR322-ColV’) using isothermal assembly. The resulting plasmids were transformed into WT or Δ*cvpA* strains.

To construct inducible N- or C-terminal sfGFP fusions to CvpA, the sfGFP sequence was amplified from pBad-sfGFP (Addgene plasmid #85482) and linked to the *cvpA* sequence amplified from EHEC genomic DNA with the linker sequence gcagcggccggcggaggg through isothermal assembly into pMMB207 (63) (ATCC #37809) multiple cloning site (MCS). Similarly, to construct the *rpoE* expression plasmid, the *rpoE* sequence was amplified from genomic DNA and cloned into the pMMB207 MCS. The resulting plasmids were transformed into WT or Δ*cvpA* strains by electroporation.

### ColV production assay

pBR322-ColV was transformed into WT or Δ*cvpA* strains. Individual colonies of either strain were selected and grown overnight in LB-Carb, and then normalized to OD 0.5. To prepare indicator plates, 100 μL of WT EHEC was added to 5 mL of 0.8% molten LB agar, vortexed to mix, then poured quickly onto a pre-warmed LB plates, which were allowed to solidify undisturbed. 5 μL of OD 0.5 ColV producing strain (WT or Δ*cvpA* transformed with pBR322-ColV) was jabbed into soft agar plate and incubated upright overnight at 37°C. The next day, plates were examined for zones of clearing around the ColV producing strain.

### DOC sensitivity assay

To determine sensitivity to DOC, each strain was grown at 37°C until mid-exponential phase (OD_600_ = 0.5). For inducible expression, at mid-exponential phase, IPTG was added to a final concentration of 1 mM and cells were incubated at 37°C shaking for 1 hour. Cultures were normalized to OD_600_ = 0.5, serially diluted and plated onto LB agar plates with and without 1% DOC and 0.1 mM IPTG.

### Microscopy

To prepare sfGFP-CvpA fusion cells for microscopy, strains were grown to mid-exponential phase (OD_600_ = 0.5). IPTG was then added to a final concentration of 1 mM. After one hour, cells were stained with the membrane dye FM4-64 as described previously (38, 64). Briefly, 1 μg/mL FM4-64 was added to 100 μL of culture and incubated at room temperature for 5 minutes. To prepare pColV-containing cells for microscopy, overnight culture was first normalized to OD_600_ = 0.5. 3 μL culture were then immobilized on 0.8% agarose pads and imaged with a Nikon Ti Eclipse microscope equipped with a widefield Andor NeoZyla camera and a 100x oil phase 31.4 numerical aperture objective. Images were processed in ImageJ/FIJI (65).

### Minimum inhibitory concentration (MIC)

To determine MIC of various compounds listed in Table 1, an assay was performed as described previously (22, 66). Briefly, compounds to be tested were prepared in serial 2-fold dilutions in LB (50 μL total volume) in a 96 well plate. Overnight cultures of bacterial strains were diluted 1:1000 in LB, grown for 1 hour at 37°C, and then diluted again 1:1000 in LB. 50 μL of this diluted culture was added to each well. Plates were incubated statically for 24 hours at 37°C.

### Transposon-insertion library construction

TIS libraries were generated in EHEC EDL933 Δ*lacI*::*aacC1* and Δ*cvpA* mutant as described previously (22). Briefly, conjugation was performed to transfer the transposon-containing suicide vector pSC189 (67) from a DAP-auxotrophic donor strain (*E. coli* MFDλpir) into the recipient strain. 600 μL of overnight culture of donor and recipient were pelleted, washed with 1 mL LB, and resuspended each in 60 μL LB. The cultures were mixed and spotted onto a 0.45 μM HA filter (Milipore) on an LB-DAP agar plant and incubated at 37°C for 1 hour. The filters were then washed in 24 mL LB and immediately spread across three 245×245 mm^2^ (Corning) LB-agar plates containing Gm and Km and incubated overnight at 37°C. Plates were scraped to collect colonies, which were resuspended in LB and stored in LB + 20% glycerol (v/v) 1 mL aliquots at −80°C. An aliquot was thawed and gDNA isolated for analysis.

### DOC-selected TIS library

1 aliquot of WT and Δ*cvpA* library were thawed and diluted to OD_600_ = 1 with LB. 5 mL of this diluted culture was added to a flask with 75 mL of LB. The culture was grown shaking at 37°C for 1 hour, at which their OD_600_ measured ~0.25. 5 mL of the culture was plated on each of two 245×245 mm^2^ (Corning) LB-agar plates containing Gm and Km and on each of two plates containing Gm and Km and 1% deoxycholate (DOC). Plates were incubated overnight (~16 hours) at 37°C and immediately scraped to collect colonies. Colonies were resuspended in LB and stored in LB + 20% glycerol (v/v) 1 mL aliquots at −80C.

### Characterization of transposon-insertion libraries

Transposon-insertion libraries were characterized as described previously (22). Briefly, for each library, gDNA was isolated using the Wizard Genomic DNA extraction kit (Promega). gDNA was fragmented to 400-600 bp using a Covaris E200 sonicator and end-repaired using the NEB Quick Blunting Kit. PCR was used to amplify transposon junctions, and PCR products were gel purified to recover 200-500 bp fragments. To estimate library concentration, purified PCR products were subjected to qPCR using primers designed to the Illumina P5 and P7 hybridization sequences. Libraries were mixed in an equimolar fashion and sequenced with a MiSeq.

Reads were trimmed of transposon and adapter sequences using CLC Genomics Workbench (Qiagen) and mapped to *E. coli* O157:H7 EDL933 (NCBI Accession Numbers: chromosome, NZ_CP008957.1; pO157 plasmid, NZ_CP008958.1) using Bowtie, allowing for no mismatches. Reads were discarded if they did not map to any sites in the genome, and reads mapping to multiple sites were randomly distributed. The data was normalized for chromosomal replication biases and differences in sequence depth using a LOESS correction of 100,000 bp (chromosome) and 10,000 bp (plasmid) windows. Reads at each TA site were tallied and binned by protein-coding gene.

For identification of loci with synthetic fitness between WT and Δ*cvpA* mutant, the Con-ARTIST pipeline was used (19, 22). First, the WT library normalized to simulate any bottlenecks or differences in sequencing depth as observed in the Δ*cvpA* library using multinomial distribution-based random sampling (n=100). Next, a Mann-Whitney U test was applied to compare these 100 simulated data sets to the Δ*cvpA* mutant library. Genes for which there were greater than 5 individual transposon-insertion mutants were considered to have sufficient data for analysis (otherwise ‘insufficient data’). Genes for which mutant abundance in the Δ*cvpA* background was mean log2FC ≥ 2 compared to the WT with a p-value <0.05 were considered to be ‘synthetic healthy’, while genes with mutant abundance log2FC ≤ −2 compared to the WT with a p-value < 0.05 were considered to be ‘synthetic sick’. The remaining genes were classified as ‘neutral’.

A similar analysis was performed to identify mutants conditionally depleted or enriched in 1% DOC compared to LB for each strain. First, for each strain, the LB library was normalized to simulate any bottlenecks or differences in sequencing depth as observed in the corresponding 1% DOC library using multinomial distribution-based random sampling (n=100). Next, a Mann-Whitney U test was applied to compare these 100 simulated data sets to the 1% DOC mutant library. Genes for which there were greater than 5 individual transposon-insertion mutants were considered to have sufficient data for analysis (otherwise ‘insufficient data’). Genes for which mutant abundance in the 1% DOC background was mean log2FC ≥ 2 compared to the WT with a p-value < 0.05 were considered to be ‘enriched’, while genes with mutant abundance log2FC ≤ −2 compared to the WT with a p-value < 0.05 were considered to be ‘depleted’. The remaining genes were classified as ‘neutral’.

### Spontaneous suppressor isolation and sequencing

To isolate and identify spontaneous suppressor mutations, the Δ*cvpA* mutant was first grown to mid-exponential phase (OD_600_ = 0.5). 2 mL of this culture was plated on 150×15mm (Fisher) LB-agar plate containing 1% DOC and incubated for 24 hours at 37°C. The next day, suppressor colonies were picked and re-streaked on LB agar plates containing 1% DOC. 15 suppressor colonies were picked, grown overnight, and subjected to gDNA isolation with the Wizard Genomic DNA extraction kit (Promega). gDNA from the Δ*cvpA* mutant was also isolated as the parental control strain. gDNA was either submitted for sequencing at the Microbial Genome Sequencing Center at the University of Pittsburgh or prepared for whole genome sequencing in house using the Nextera XT DNA Library Preparation Kit (Illumina). Libraries were sequenced using the NextSeq 550 or MiSeq platforms, respectively.

Reads were mapped to the *E. coli* O157:H7 EDL933 genome (NCBI Accession Numbers: chromosome, NZ_CP008957.1; pO157 plasmid, NZ_CP008958.1) using CLC Genomics Workbench (Qiagen) with default parameters. Basic variant detection was performed, assuming a ploidy of one and filtering out variants with a frequency of less than 35%. Known variants were filtered against the Δ*cvpA* strain. Variant calls were then manually curated to identify SNPs with a frequency greater than 90%.

### Bioinformatics / protein prediction

Protein predictions about CvpA structure were inferred from several programs: PSLPred, HHPred, Phobius, Phyre2, and OCTOPUS (68–72). Protter (73) was used to create a topological diagram. This figure originally appeared in (22).

The topology diagram for CvpA was made using Protter. To examine the conservation of CvpA amino acid sequence across bacterial phyla, we used Annotree version 89 (50). Query term ‘CvpA’ (KEGG K03558) was used to find bacterial proteins with at least 30% identity and 70% alignment across 27,000 genomes. 20 CvpA sequences from genomes across bacterial phyla were selected for comparison and aligned using MUSCLE (74) within the MEGA X platform (75) using default parameters.

## Acknowledgements/Funding

We are grateful to members of the Waldor lab, particularly Jacob Lazarus, for comments on this project and the manuscript. MKW is funded by NIH grant R01-AI-042347 and the Howard Hughes Medical Institute.

## Supplemental Figures

**Figure S1:**
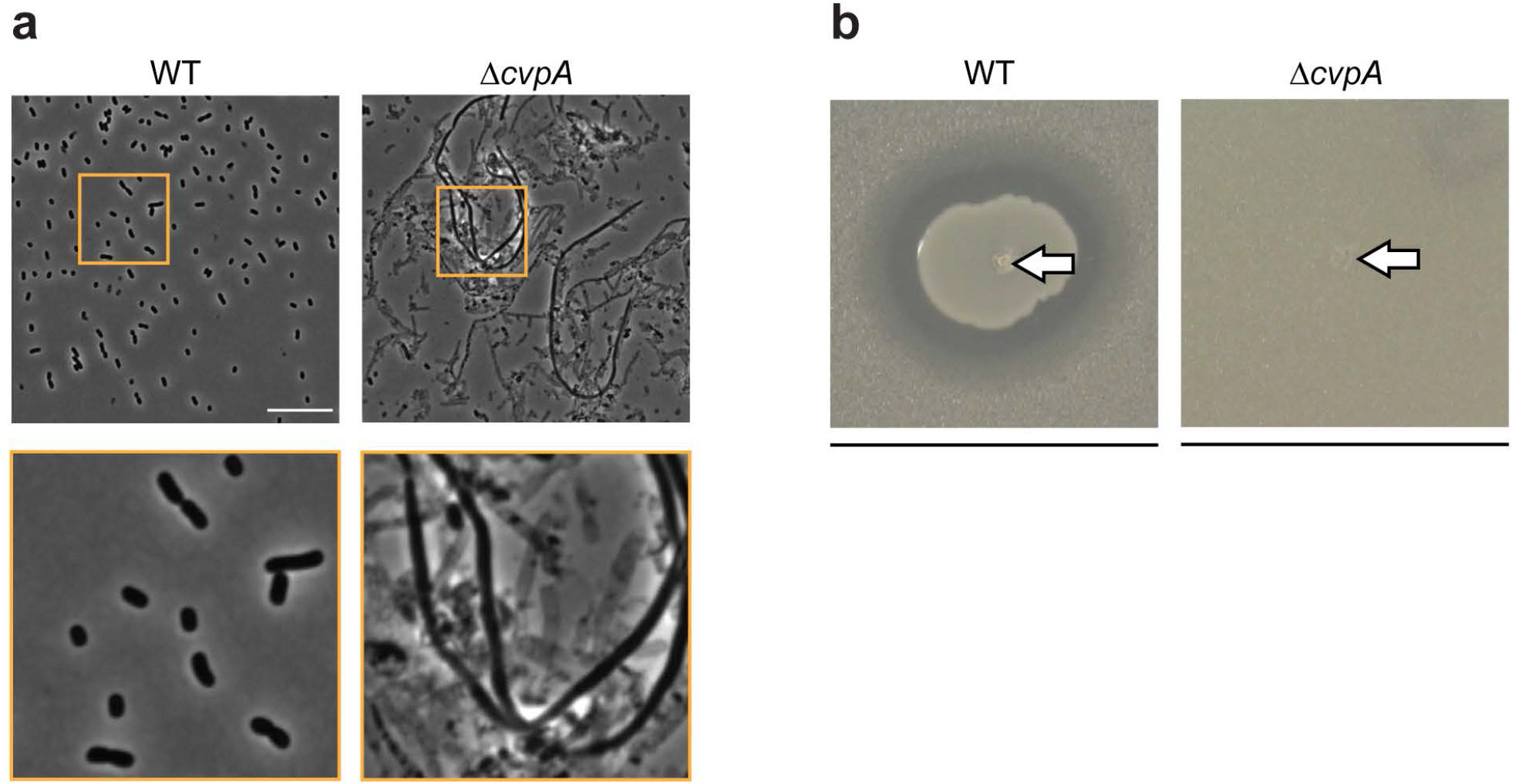
Δ*cvpA* EHEC is sensitive to Colicin V production and does not appear to secrete it. a) Micrograph of overnight culture of indicated strains expressing ColV from plasmid pBR322-ColV. Scale bar is 10 μM. Orange box denotes inset. b) 5 uL jab of overnight culture of indicated strain in soft agar matrix containing 100 μL of indicator (sensitive EHEC) strain. Arrow indicates position of jab. Line represents 20 mm.

**Figure S2:**
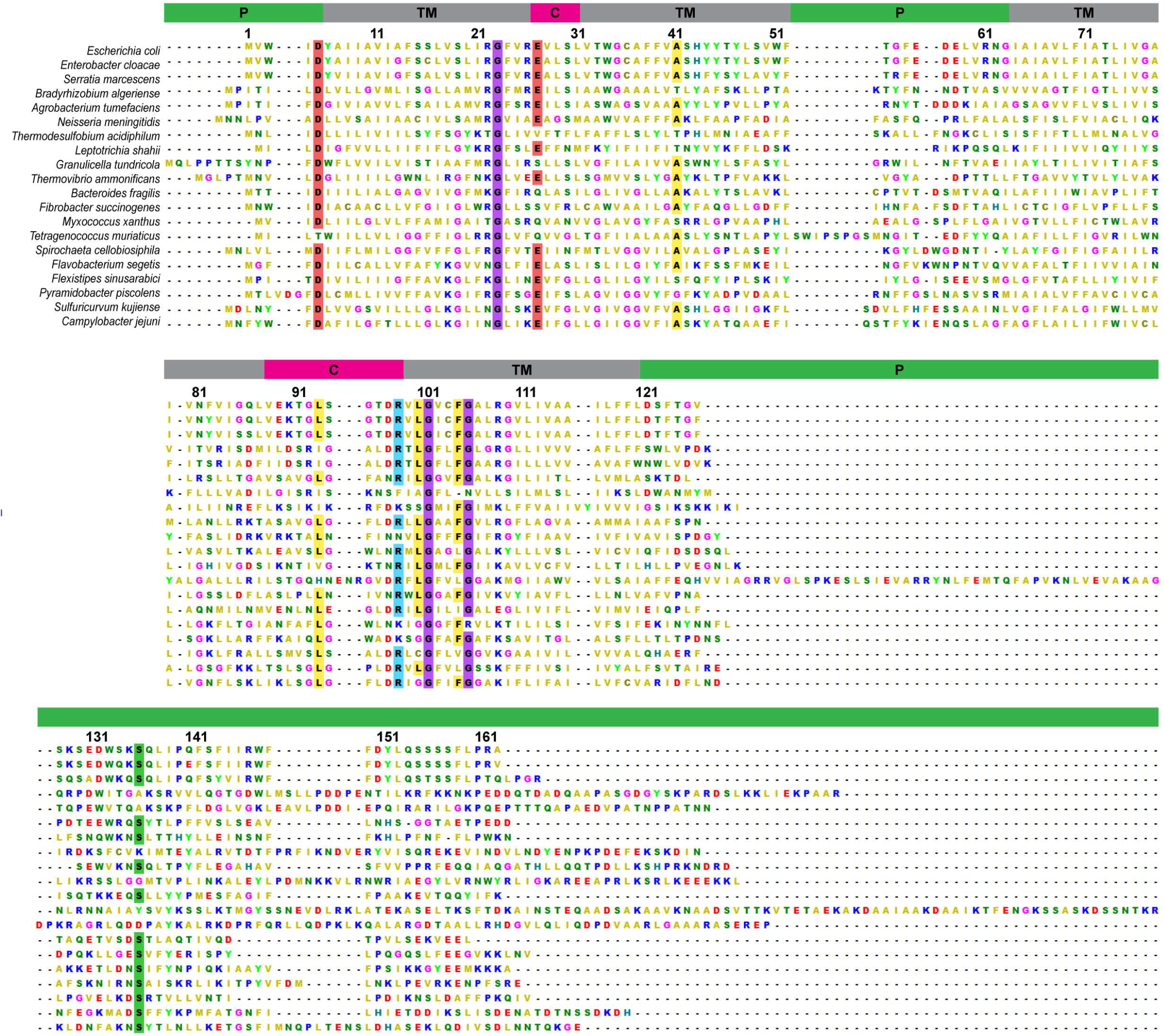
MUSCLE alignment of 20 CvpA homologs from bacterial species across diverse phyla. Residues with >70% conservation across species are indicated by a colored box. Residue numbering and predicted membrane topology for the *E. coli* CvpA is shown above the alignment. The green box indicates predicted periplasmic (P) segments, the gray box indicates transmembrane (TM) segments, and the pink box represents cytoplasmic (C) segments.

